# SARS-CoV-2 Delta Spike Protein Enhances the Viral Fusogenicity and Inflammatory Cytokine Production

**DOI:** 10.1101/2021.11.23.469765

**Authors:** Zhujun Ao, Maggie Jing Ouyang, Titus Abiola Olukitibi, Xiaojian Yao

**Affiliations:** Laboratory of Molecular Human Retrovirology, Department of Medical Microbiology, Max Rady College of Medicine, Rady Faculty of Health Sciences, University of Manitoba, Winnipeg, MB, Canada

**Keywords:** COVID-19, SARS-CoV-2, Delta variant, Spike protein, cell-to-cell fusion, NF-κB pathway, proinflammatory cytokines

## Abstract

The Delta variant is now the most dominant and virulent SARS-CoV-2 variant of concern (VOC). In this study, we investigated several virological features of Delta spike protein (SP_Delta_), including protein maturation and its impact on viral entry of cell-free pseudotyped virus, cell-cell fusion ability and its induction of inflammatory cytokine production in human macrophages and dendritic cells. The results showed that SPΔC_Delta_ exhibited enhanced S1/S2 cleavage in cells and pseudotyped virus-like particles (PVLPs). We further showed that SPΔC_Delta_ elevated pseudovirus infection in human lung cell lines and mediated significantly enhanced syncytia formation. Furthermore, we revealed that SPΔC_Delta_-PVLPs had stronger effects on stimulating NF-κB and AP-1 signaling in human monocytic THP1 cells and induced significantly higher levels of pro-inflammatory cytokine, such as TNF-α, IL-1β and IL-6, released from human macrophages and dendritic cells. Overall, these studies provide evidence to support the important role of SPΔC_Delta_ during virus infection, transmission and pathogenesis.

## INTRODUCTION

The emergence of the highly pathogenic coronavirus disease 2019 (COVID-19) has been a major concern and threat to public health for two years. As of early November 2021, approximately 248 million COVID-19 cases and more than 5 million deaths were reported globally (WHO, 2021). COVID-19 is caused by severe acute respiratory syndrome coronavirus 2 (SARS-CoV-2), which is a member of betacoronaviridae with a single-stranded 30 kb positive-sense RNA genome encoding 29 proteins (Srivastava et al., 2021). Within 2 years, multiple variants of SARS-CoV-2 have emerged (Galloway et al., 2021; Greaney et al., 2021; Nonaka et al., 2021; Paiva et al., 2021; Resende et al., 2021; Santos and Passos, 2021; Tegally et al., 2020; Volz et al., 2021). Delta variant, which was first detected in India and derived from the Pango lineage B.1.617.2, is the most dominant variant of concern (VOC) and has accounted for approximately 99% of new cases of coronavirus worldwide (Banu et al., 2020; Ranjan et al., 2021; Sahoo et al., 2021; Worldometer, 2021). Previous studies suggested that Delta variant infection has a shorter incubation period but a greater viral load (> 1,000 times) than earlier variants (Li et al., 2021; Reardon, 2021). Further, the patients contracted with Delta variant have higher hospitalization rate and more severe outcomes (Twohig et al., 2021). Unfortunately, current vaccines only provide partial protection against the infection of Delta variant, because these vaccines are designed based on the original Wuhan-Hu-1 sequence (Mlcochova et al., 2021). Therefore, it is important to understand the molecular mechanisms of the increased transmissibility and immune evasion of these SARS-CoV-2 variants to facilitate the development of vaccines and therapeutic drugs against COVID-19.

VOCs are mainly classified based on the mutations on their spike protein (SP). The SP of coronavirus is responsible for viral attachment and entry to the host cells (Huang et al., 2020b). The matured SP is cleaved to generate S1 and S2 subunits at specific cleavage sites. The S1 subunit (aa 14-685) is responsible for receptor binding through its receptor-binding domain (RBD). The S2 subunit (aa 686-1273) mediates membrane fusion to facilitate cell entry (Bertram et al., 2013; Hoffmann et al., 2020b; Peacock et al., 2021b). Mutations in SP have resulted in the high rates of transmission and replication of various variants (Zhang et al., 2021a; Zhou et al., 2021) and the immune evasion from antibody neutralization of various variants (McCallum et al.; Mlcochova et al., 2021; Zhang et al.). The SP of the Delta variant has eight mutations compared with the original virus, including T19R, Δ156– Δ157, and R158G in the N-terminus Domain (NTD), D614G, L452R and T478K at the RBD, P681R close to the furin cleavage site, and D950N at the S2 region (Cherian et al., 2021; Planas et al., 2021; Zhang et al., 2021a). It has been demonstrated that the mutations at the NTD of Delta SP alter the antigenic surface near the NTD-1 epitope, thus leading to the lack of binding affinity with the NTD neutralizing antibodies (Zhang et al., 2021a).

Furthermore, the P681R mutation closed to the furin cleavage has been demonstrated to aid the pathogenicity of the virus (Cherian et al., 2021; Liu et al., 2021; Saito et al., 2021). The newly identified furin cleavage site (681-PRRAR↓SV-687) at the S1/S2 site is reported to be critical for the pathogenesis of SARS-CoV-2 in mouse models, and also be responsible for the cell-cell fusion, which is absent in other group-2B coronaviruses (Coutard et al., 2020; Johnson et al., 2020; Xia et al., 2020). It was shown that P681R mutation at this cleavage site endow Delta variant with special features that facilitate the spike protein cleavage and viral fusogenicity (Peacock et al., 2021b; Saito et al., 2021). Study found that a chimeric Delta SARS-CoV-2 bearing the Alpha-SP replicated less efficiently than the wild-type Delta variant, and the reversion of Delta P681R mutation to wild-type P681 attenuated Delta variant replication as well (Liu et al., 2021). These observations suggested that the P618R mutation that occurs in Delta variants contributes immensely to the high replication and transmissibility rate. We are therefore interested in further investigating how the P618R mutation impacts the high replication rate of Delta variants.

Like other high pathogenic viruses (influenza H5N1, SARS-CoV-1 and MERS-CoV), SARS-CoV-2 infection also induced excessive inflammatory response with the release of a large amount of pro-inflammatory cytokines (cytokine storm) that may result in Acute Respiratory Distress Syndrome (ARDS) and multiorgan damage (Zhu and al., 2020). Clinical studies showed that the high mortality of COVID-19 is related to cytokine release syndrome (CRS) in a subgroup of severe patients (Huang et al., 2020a), characterized by elevated levels of certain cytokines including IL-6, TNF-α, IL-8, IL-1β, IL-10, MCP-1 and IP-10 (Hadjadj et al., 2020; Huang et al., 2020a; Mehta et al., 2020; Xu et al., 2020; Yang et al., 2020). The observed cytokine production induced by SARS-CoV-2 infection or spike protein expression has been linked with nuclear factor kappa B (NF-κB) and and activator protein-1(AP-1) signaling pathways that can induce the expression of a variety of proinflammatory cytokine genes (Neufeldt et al., 2020b; Zhu et al., 2021). Like other RNA virus, SARS-CoV-2 was found activate NF-κB and AP-1 transcription factors following the sensing of viral RNAs or proteins by different pathogen pattern recognition receptors (PRRs) and associated signalling cascades, including RLRs and TLRs (Pantazi et al., 2021; Zhu et al., 2021). In addition, angiotensin II type 1 receptor (AT1)-MAPK signalling (Patra et al., 2020) and the cGAS-STING signalling (Neufeldt et al., 2020a) have been identified responsible for the activation of NF-κB and the elevated expression of IL-6 in the SARS-CoV-2 infected or SP expressing cells. However, whether Delta variant or its SP may initiate stronger cytokine storm in patients that leading to more severe illness than other strains still required more investigation.

The current study aims to characterize the cleavage/maturation of various SPs of SARS-CoV-2, especially the Delta variant SP, and their effects on virus infection, cell-cell fusion, cytokine production and related signaling pathways. By using a SARS-CoV-2 SP pseudotyped lentiviral vector or viral-like particles (PVLPs), we demonstrated that SP_Delta_ enhanced S1/S2 cleavage, accelerated pseudovirus infection and promoted cell-cell fusion. We also showed that SP_Delta_ strongly activates NF-κB and AP-1 signaling in THP1 cells. Furthermore, we observed that SPΔC_Delta-_PVLP stimulation promoted the production of several pro-inflammatory cytokines by human macrophages and dendritic cells.

## RESULTS

### SARS-CoV-2 Delta SP exhibited enhanced cleavage and maturation in cells and in the pseudotyped virus

To investigate the functional role of SARS-CoV-2 Delta SP, we first synthesized cDNA encoding SARS-CoV-2 Delta SP, as described in CDC’s SARS-CoV-2 Variant Classifications and Definitions (CDC, 2021) (Fig. 1A) and inserted cDNA into a pCAGG-expressing plasmid, as described in the Materials and Methods. Previously described pCAGG-SPΔC_WT-_ and pCAGG-SPΔC_G614_-expressing plasmids (Ao et al., 2021a) were also used in this study. Meanwhile, we constructed a pCAGG-SPΔC_Delta-PD_ in which arginine (R) at the amino acid position of 681 and asparagine (N) at 950 in SPΔC_Delta_ were reverted to the original proline (P) and aspartic acid (D) to test the effect of these amino acids on the function of SPΔC_Delta_ (Fig. 1A). To enhance the transportation of SP to the cell surface and increase the virus incorporation of SP, the cDNA encoding the C-terminal 17 aa in SARS-CoV-2 SP was deleted in all pCAGG-SPΔC plasmids (Ao et al., 2021a).

**Figure 1.**
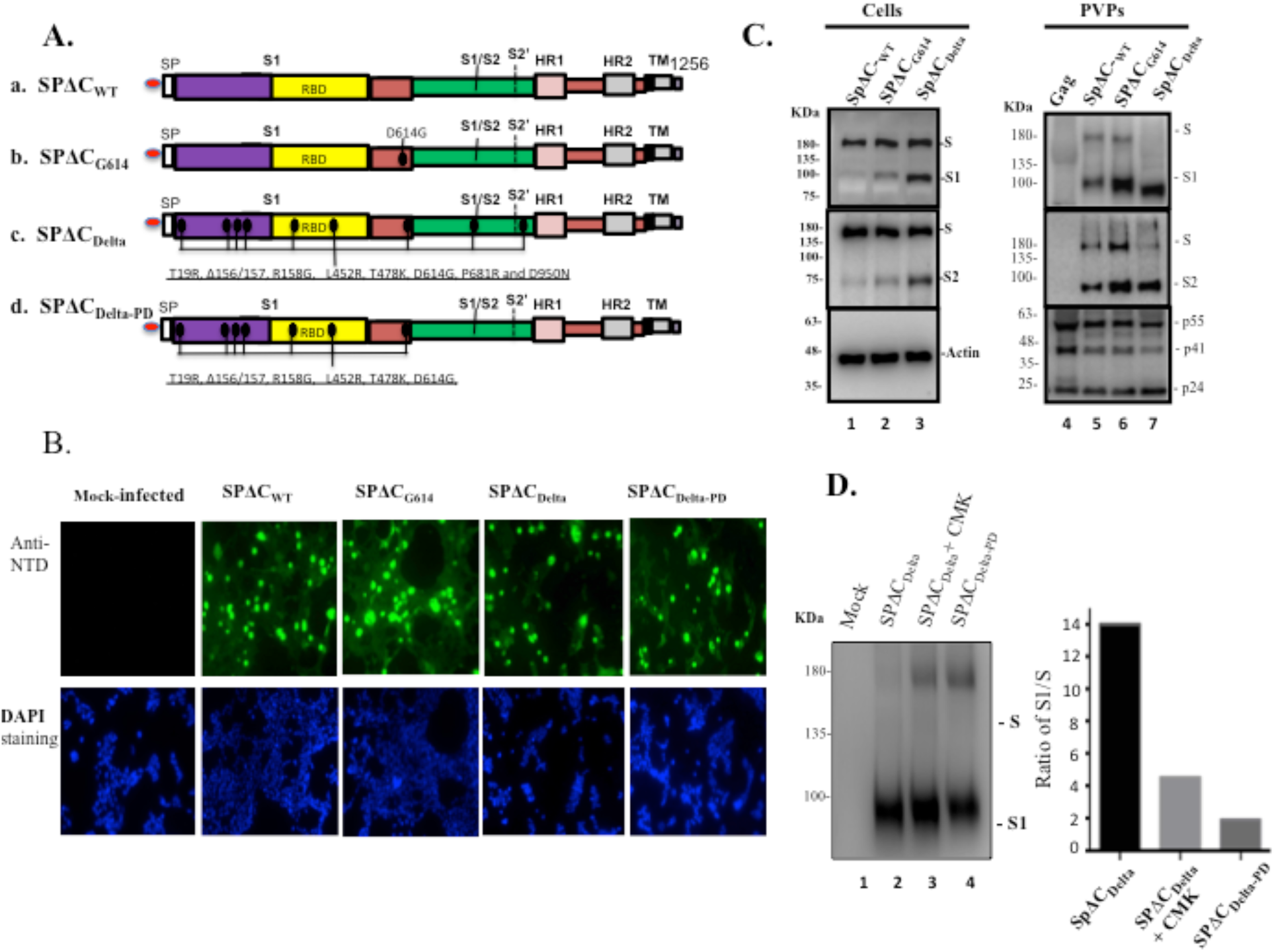
The expression and cleavage of SARS-CoV-2 Delta Spike protein (SP) in the cells and in the pseudotyped virus particles (PVPs). (A) Schematic representation of the structure domains and mutations of SARS-CoV-2 SP and its variants. A C-terminal 17aa of each of SARS-CoV-2 SPΔC was deleted in order to increase the incorporation of SP into the virrus particles. (B) 293T cells were transfected with pCAGGS-SPΔC_wt_, -SPΔC_G614_, -SPΔC_Delta_ or - SPΔC_DeltaPD_. The intracellular expression of each SPΔC in the transfected 293T cells were detected by immunofluorescence assay with SARS-CoV-2 S-NTD antibody. (C) 293T cells were transfected with each SPΔC-expressing plasmid, HIV ΔRI/Env^-^/Gluc) and a packaging plasmid (pCMVΔ8.2) in HEK293T cells. As Gag-control, 293T cells were only transfected with pCMVΔ8.2 plasmid. WB were used to detect the expression and cleavage of various SPΔC and HIV Gag protein in cells and in PVPs. Full-length spike (S), cleaved S1 and S2 were annotated. (D) SPΔC_Delta-_PVPs were produced in the absence or in the presence of a furin inhibitor (CMK) (25μM) and SPΔC_DeltaPD_-PVPs were analyzed by WB using polyclonal anti-SP/RBD antibody (Left panel). The ratio of S1 relative to the full length SP was also quantified by laser densitometry as indication of SP processing efficiencies (Right panel). Panel C-D show the representative WB image from three independent experiments.

To examine the expression of various SPs in the cells and their incorporation in the SPΔC-pseudotyped viral particles (PVPs), each SPΔC-expressing plasmid was cotransfected with a multiple-gene deleted HIV-based vector encoding a Gaussia luciferase gene (ΔRI/ΔEnv/Gluc) and a packaging plasmid (pCMVΔR8.2) in HEK293T cells, as described previously (Ao et al., 2021a). After 48 hrs of transfection, the expression of each SPΔC in the transfected cells was analyzed by an indirect immunofluorescence (IF) assay using human SARS-CoV-2 S-NTD antibodies. The results revealed that all SPΔCs were well expressed in the transfected cells or cell surface (Fig. 1B). Meanwhile, the transfected cells and PVPs were lysed and processed with WB with anti-SP/RBD or anti-S2 antibodies, respectively (Fig. 1C, top and middle panels). Interestingly, the data clearly showed that in the cells, SPΔC_Delta_ was more efficiently processed from the S precursor into S1 and S2 than SPΔC_WT_ and SPΔC_G614_ (Fig. 1C, compare Lane 3 to Lanes 1 and 2). In PVPs, the majority of SPΔC_Delta_ presented as a mature form (S1 and S2) compared to SPΔC_WT_ and SPΔC_G614_ (Fig. 1C, compare Lane 7 to Lanes 5 and 6), indicating that SPΔC_Delta_ undergoes a more efficient maturation process. Surprisingly, S1 of SPΔC_Delta_ but not S2 appeared to migrate faster than S1 of SPΔC_WT_ and SPΔC_G614_ (Fig. 1C, top panel). The possible mechanism for this behavior is currently unknown.

The cleavage of the SARS-CoV-2 SP into S1 and S2 most likely occurs by furin, and the P681R mutation of the SP Delta was suggested to enhance S1/S2 cleavability (Liu et al., 2021; Peacock et al., 2021a). We therefore further tested whether a fusin protease inhibitor, a peptidyl chloromethylketone (CMK), or the reverse change in R_681_ of SPΔC_Delta_ to P would alter the maturation rate of the S protein. The SPΔC_Delta-PD-_PVPs or SPΔC_Delta_-PVPs packaged in the presence or absence of CMK were analyzed by WB with anti-RBD antibody and quantified by densitometry using ImageJ (https://imagej.nih.gov/ij/). The results showed that either CMK treatment or SPΔC_Delta-PD_ clearly negatively impacted the maturation of SPΔC_Delta_ (Fig. 1D).

### Delta-SP mediates more efficient pseudovirus infection in a lung epithelial cell line and primary macrophages

To investigate the impact of SPΔC_Delta_ on viral infection, we produced Gluc-expressing Delta-SP-PVPs (Fig. 2A) and tested the infectivity of different pseudoviruses in two human lung epithelial cell lines, Calu-3 and A549 cells. To increase susceptibility to SP-PVP infection, the A549_ACE2_ cell line was generated by transducting a lentivirus expressing hACE2 and subsequent puromycin selection, as described in the Materials and Methods. hACE2 expression in A549_ACE2_ cells was verified by WB (Fig. 2B). Then, both cell lines were infected with equivalent amounts (adjusted with p24 values) of the SPΔC_WT_-, SPΔC_G614_-, and SPΔC_Delta_-PVPs for three hours and washed. At 24 and 48 hrs post infection (p.i.), the supernatants were collected, and the infection levels of pseudoviruses were monitored by measuring Gluc activity. The results showed that in both cell lines, the SPΔC_Delta_-PVPs exhibited the highest infection efficiency, the SPΔC_G614_-PVPs had a slightly lower infection efficiency than SPΔC_Delta_, while the SPΔC_wt_-PVPs showed a significantly lower infection efficiency (Fig. 2C). All of these results indicated that SPΔC_Delta_-PVPs had a significantly more efficient virus entry step than SPΔC_wt._-PVPs in a single cycle replication system.

**Figure 2.**
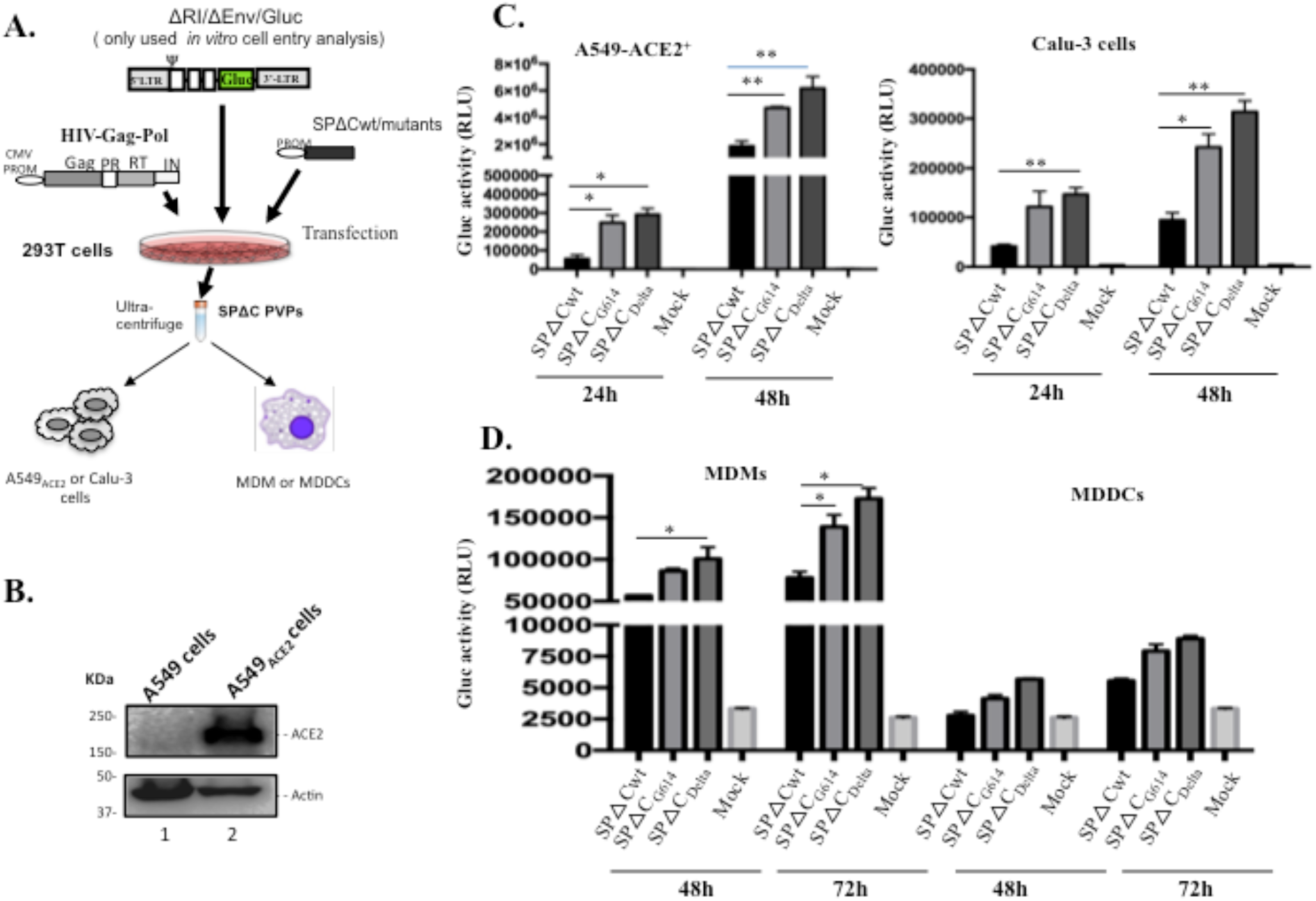
SP pseudotyped viruse infectivity assays on human lung cell lines, human macrophages (MDMs) and dendritic cells (MDDCs). (A) Schematic representation of the procedures and the plasmids used for production of SARS-COV2-SPΔC-pseudotyped lentivirus particles (SPΔC-PVPs). (B) an A549_ACE2_ cell lins was generated by transducing A549 cells with a lentiviral vector (pLenti-C-ACE2).The ACE2 expression in the transduced cells was detected by WB using anti-ACE2 antibody. (C, D) A549_ACE2_, Calu-3 cell lines, human MDMs or MDDCs were infected with an equal amounts of SPΔC_wt_, -SPΔC_G614_, or - SPΔC_Delta_-PVPs carrying Gaussia luciferase (Gluc) gene (adjusted by P24). At different time points, the Gluc activity in the supernatant of infected cultures was measured. The results are the mean values ± standard deviations (SD) of two independent experiments. Statistically significant differences (* P ≤0.05; **, P ≤ 0.01) versus the SPΔCwt were determined by unpaired t test. No significant (ns) was not shown.

Next, we also checked the ability of SPΔC-PVPs to infect human differentiated macrophages and dendritic cells. Briefly, human monocyte-derived macrophages (MDMs) or dendritic cells (MDDCs) were infected with equal amounts (adjusted with HIV p24 levels) of SPΔC_wt-_-, SPΔC_G614_-, and SPΔC_Delta_-PVPs. At 48 and 72 hrs p.i., the Gluc activity in the supernatant from the infected cell cultures was monitored. The results showed that both human primary cells, especially MDMs, could be infected by SPΔC-PVPs, while SPΔC_Delta_- and SPΔC_G614_-PVPs displayed more efficient infection than SPΔC_wt_-PVPs (Fig. 2D). All of these experimental observations indicate that SPΔC_Delta_-PVPs have a stronger ability to target MDMs than SPΔC_wt_-PVPs. The results also suggested that MDDCs can be targeted by SPΔC-PVPs but with less efficiency.

### Delta-SP variant enhanced syncytia formation in lung epithelial A549 cells expressing ACE2

Previous studies have shown that SARS-CoV-2 SP is able to possess fusogenic activity and form large multinucleated cells (syncytia formation) (Bussania et al., 2020; Cattin-Ortolá et al., 2020). We then asked whether Delta-SP could possess higher fusogenic activity than other variants. Briefly, 293T cells were transfected with SPΔC_WT_, SPΔC_G614_, SPΔC_Delta,_ or SPΔC_DeltaPD_ plasmids by Lipofectamine 2000. At 24 hrs of transfection, we mixed SPΔC-expressing 293T cells with A549_ACE2_ cells at a ratio of 1:3. At 6 and 30 hrs post transfection, syncytial formation was observed under a microscope, and the results revealed that SPΔC_WT_ and SPΔC_G614_ induced similar levels and sizes of syncytia. Intriguingly, an increasing number of syncytia formations were observed in the coculture of SPΔC_Delta_-expressing 293T and A549_ACE2_ cells (Fig. 3A and B), indicating that SP from the Delta variant has a stronger fusogenic ability. However, SPΔC_DeltaPD_-expressing 293T/A549_ACE2_ cell coculture displayed less syncytia formation than SPΔC_Delta_ (Fig. 3 B). A and B), suggesting the importance of P681R for the strong fusogenic activity of SP from the Delta variant.

**Figure 3.**
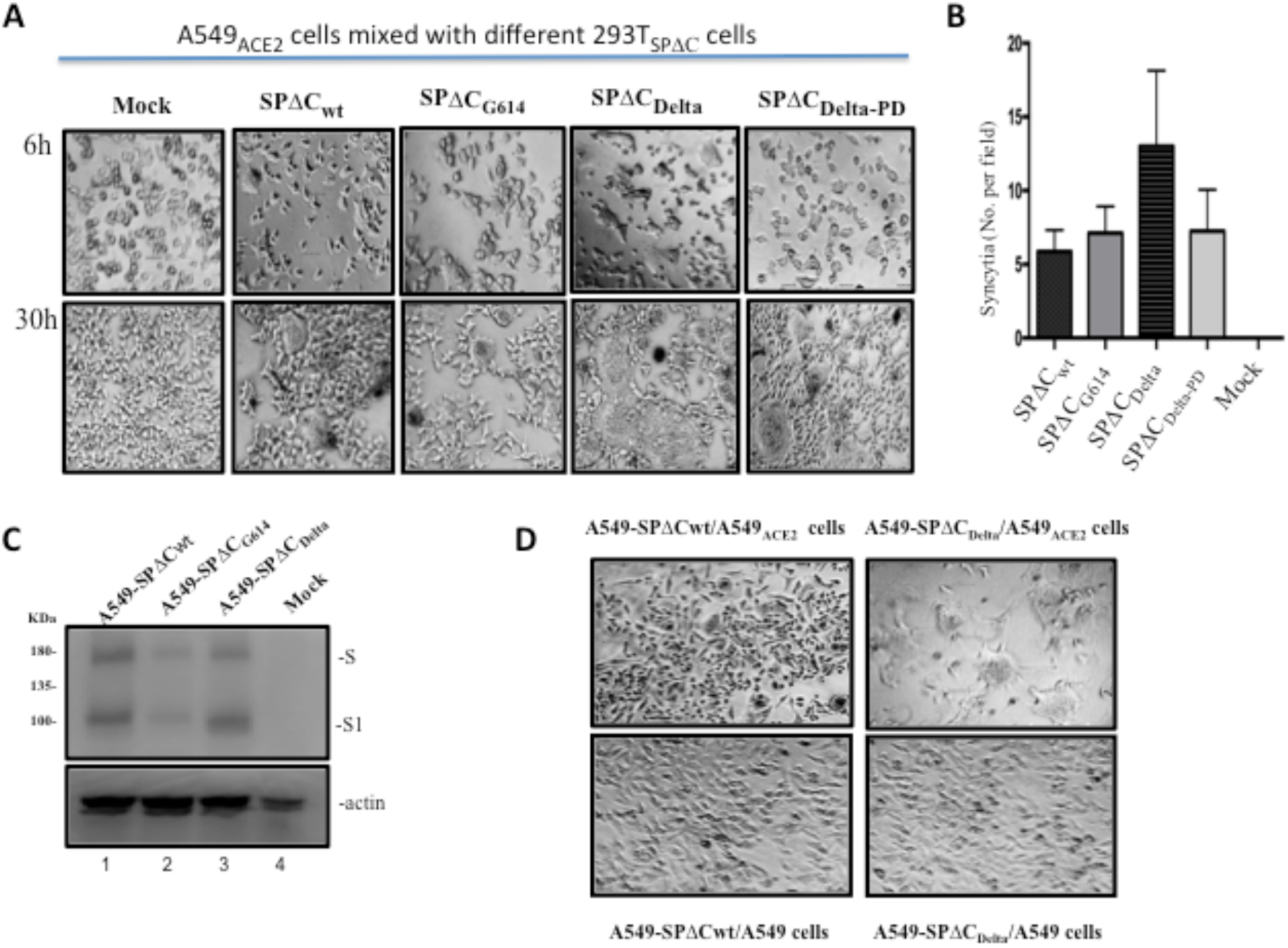
Delta-SP variant enhanced syncytia formation in lung epithelia cell line. (A) A549_ACE2_ cells were cocultured with 293T cells transfected with SPΔC or variants. The syncytia formation was monitored by microscopy at 6 hours and 30 hours after coculture. (B) Quantification of the numbers of syncytia formation in the cocultures at 6 hrs under bright-field microscopy. Results are the mean values ± standard deviations (SD) from two independent experiments. C) Expression of SPΔC_wt_, SPΔC_G614_, or SPΔC_Delta_ in corresponding A549 stably cell lines was detected by WB using anti-SP/RBD antibody. D) A549_ACE2_ cells or A549 cells were cocultured with A549-SPΔC_wt_ or A549-SPΔC_Delta_ stable cells. The syncytia formation was visualized by microscopy at 30 hours after co-culture.

To further confirm the strong fusogenic ability of SPΔC_Delta_, we also generated A549 cells stably expressing SPΔC_wt_, SPΔC_G614_ or SPΔC_Delta_ (named A549-SPΔC_Delta,_ A549-SPΔC_G614_ or A549-SPΔC_wt_ cells) (Fig. 3C). Since A549-SPΔC_wt_ and A549-SPΔC_Delta_ cells displayed similar levels of SPΔC expression based on a WB analysis, we then tested their fusogenic ability by mixing A549-SPΔC_Delta_ or A549-SPΔC_wt_ cells with the A549_ACE2_ cell line using a similar experimental process as described above. Meanwhile, A549-SPΔC_Delta_ or A549-SPΔC_wt_ cells were cocultured with A549 cells as a control. The results confirmed that the coculture of A549-SPΔC_Delta_ cells and A549_ACE2_ cells formed large syncytia formation more efficiently than that of A549-SPΔC_wt_ cells (Fig. 3D), confirming the stronger fusogenic activity of the SP of the Delta variant.

### Delta variant SP stimulates higher NFκB and AP1 signaling pathway activities

The severity of COVID-19 is highly correlated with dysregulated and excessive release of proinflammatory cytokines (Huang et al., 2020a). Given that the NFκB and AP1 signaling pathways are among the critical pathways responsible for the expression of proinflammatory cytokines and chemokines (Hojyo et al., 2020; Kawasaki and Kawai, 2014), we therefore examined the activities of these two signaling pathways triggered by SPΔC in the monocyte cell line THP1 and THP1-derived macrophages. First, we generated THP1-NF-κB-Luc and THP1-AP-1-Luc sensor cell lines by transducing THP1 cells with a lentiviral vector encoding the luciferase reporter gene driven by NFκB- or AP1-activated transcription response elements (Fig. 4A), as described in the Materials and Methods. To obtain THP1-derived macrophages, THP-1-NF-κB-Luc and THP1-AP-1-Luc sensor cell lines were treated with phorbol 12-myristate 13-acetate (PMA) (100 nM) for 3 days. Additionally, we produced genome-free SPΔC-PVLPs by cotransfecting each SPΔC-expressing plasmid with a packaging plasmid (pCMVΔR8.2) in 293T cells, and the expression of SPΔC in the purified PVLPs was verified by WB with an anti-RBD antibody (Fig. 4B). Then, different THP1 sensor cell lines and THP1-derived macrophages were treated with different SPΔC-PVLPs of same amount (adjusted by p24) for 6 hrs, and the luciferase activity in treated cells was measured by a luciferase assay system (Promega). Interestingly, we found that the NFκB activity induced by SPΔC_wt_ and SPΔC_G614_-PVLPs was slightly higher than that induced by the VLP control (Gag). However, the SPΔC_Delta_-treated THP1 cells/macrophages produced significantly higher (3∼7-fold) NFκB activity compared with SPΔC_wt_ or SPΔC_G614_ (Fig. 4C). Consistent with this finding, SPΔC_Delta_ also triggered higher (∼2-fold) AP1 signaling pathway activities in THP1/macrophages than SPΔC_wt_ or SPΔC_G614_ (Fig. 4D). These results indicated that SPΔC_Delta_ triggered significantly stronger signals to activate the NFκB and AP1 pathways in the monocyte cell line THP1 and THP1-derived macrophages.

**Figure 4.**
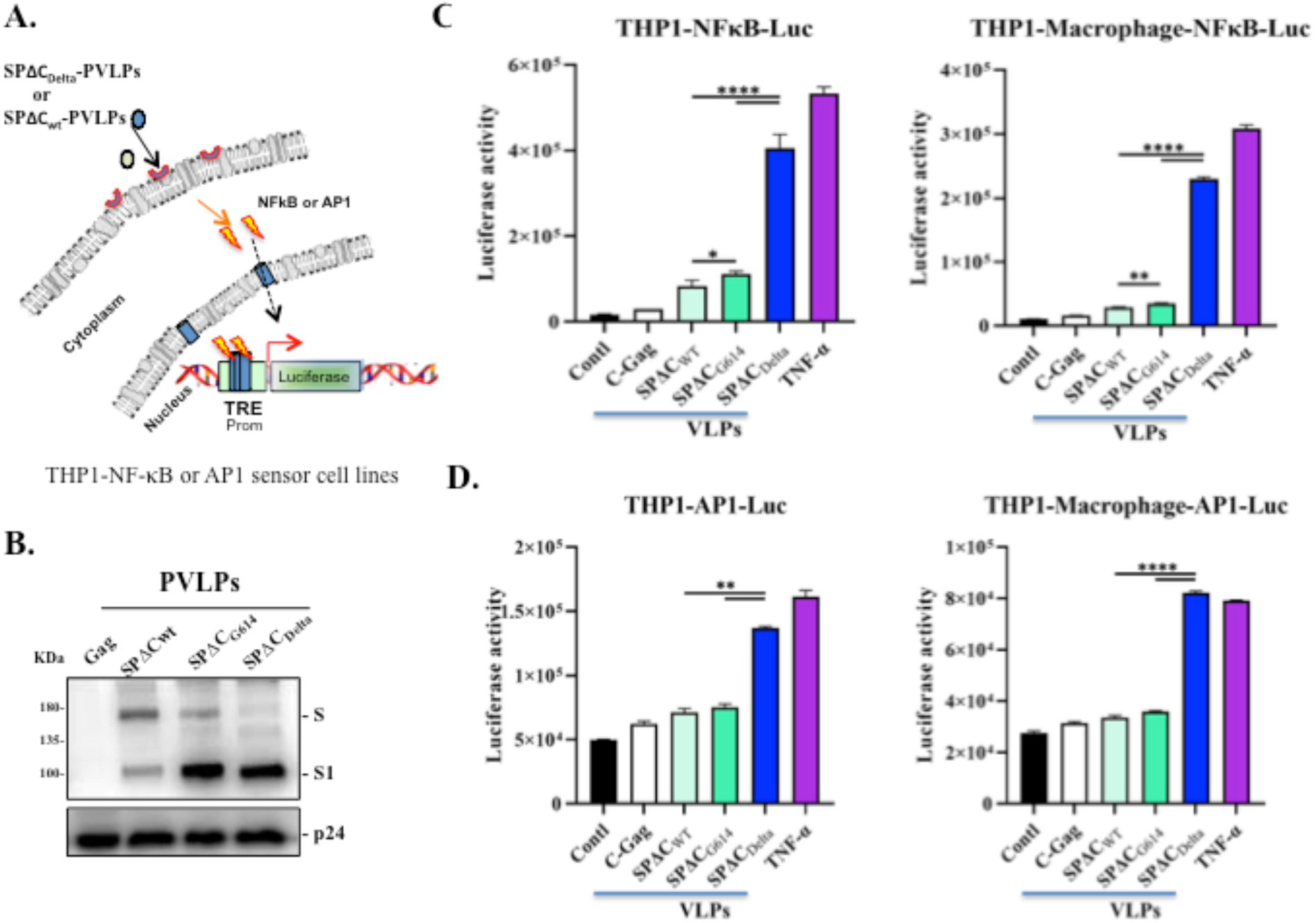
SPΔC_Delta_-PVLPs stimulated NF-κB and Ap1-signal pathway in THP1 cells and THP1 derived macrophages. **(A)** The schematic diagram of NF-κB activity luciferase reporter assay. The THP1-NF-κB -Luc, or THP1-AP1-Luc sensor cell line were incubated with SPΔC_wt_, -SPΔC_G614_, or -SPΔC_Delta_-PVLPs for 6 hrs, and the activation of NF-κB or AP1 signaling was detected by measurement of the luciferase activity. (B) WB detected the incorporation of SPΔC_wt_, -SPΔC_G614_, or -SPΔC_Delta_ in PVLPs using SARS-CoV-2 S-NTD antibody. The GagP24 was detected using mouse anti-P24 antibody. (C, D) The THP1-NFκB-Luc or THP1-NFκB-Luc-derived macrophages, and THP1-AP-1-Luc cells or THP1-AP-1-Luc-derived macrophages were treated with equal amounts of SPΔC_wt_, SPΔC_G614_ or SPΔC_Delta_-PVLPs (adjusted by P24) for 6 hrs, and the activation of NF-κB or AP-1 signaling was detected by measurement of the Luc activity. Meanwhile, Gag-VLPs or TNF-α treatment were used as negative or positive control. The results are the mean values ± standard deviations (SD) of two independent experiments. Statistical significance was determined using unpaired t test, and significant p values are represented with asterisks (*P ≤0.05; **P ≤ 0.01; ***P≤0.001; **** P≤0.0001). No significant (ns) was not shown

### Delta variant SP stimulates higher proinflammatory cytokine production in human macrophages (MDMs) and dendritic cells (MDDCs)

Previous studies have shown that SARS-CoV-2 infection can stimulate the production of immunoregulatory cytokines (IL-6, IL-10) in human monocytes and macrophages (Boumaza et al., 2021). We further investigated whether SP of the Delta variant can induce higher levels of proinflammatory cytokine and chemokine in MDMs and MDDCs. Briefly, human MDMs and MDDCs were treated with the same amount (adjusted by p24) of SPΔC-PVLPs, including SPΔC_wt_-, SPΔC_G614_-, SPΔC_Delta_-PVLPs or control VLPs (Gag-VLPs). After 24 hrs of incubation, the cytokines released in the supernatants were determined by a MSD (Meso Scale Discovery) immunoassay. The results revealed that SPΔC_wt_-PVLP stimulation did not result in a significant change in cytokine release from MDMs compared to the control VLPs (Fig. 5A). However, in MDMs, SPΔC_Delta_-PVLPs induced significantly higher levels of several proinflammatory cytokines, such as IFN-γ, TNF-α, IL-1β, and IL-6, while SPΔC_G614_-PVLPs also induced increase of these cytokines but overall to a less extent, when compared with SPΔC_wt_-PVLPs (Fig. 5A). For example, the SPΔC_Delta_-PVLPs elevated TNF-α level 61-fold in comparison with SPΔC_wt_-PVLPs, contrastingly, SPΔC_G614_-PVLPs only increased to approximately 33-fold. Nevertheless, all of SPΔC-PVLPs showed no stimulating effects on IL-2 and IL-8 production. Interestingly, SPΔC_Delta_-PVLPs and SPΔC_G614_-PVLPs also slightly increased anti-inflammatory cytokines IL-4, IL-10 and IL-13, indicating the negative feedback of inflammation may exist during these stimulations.

**Figure 5.**
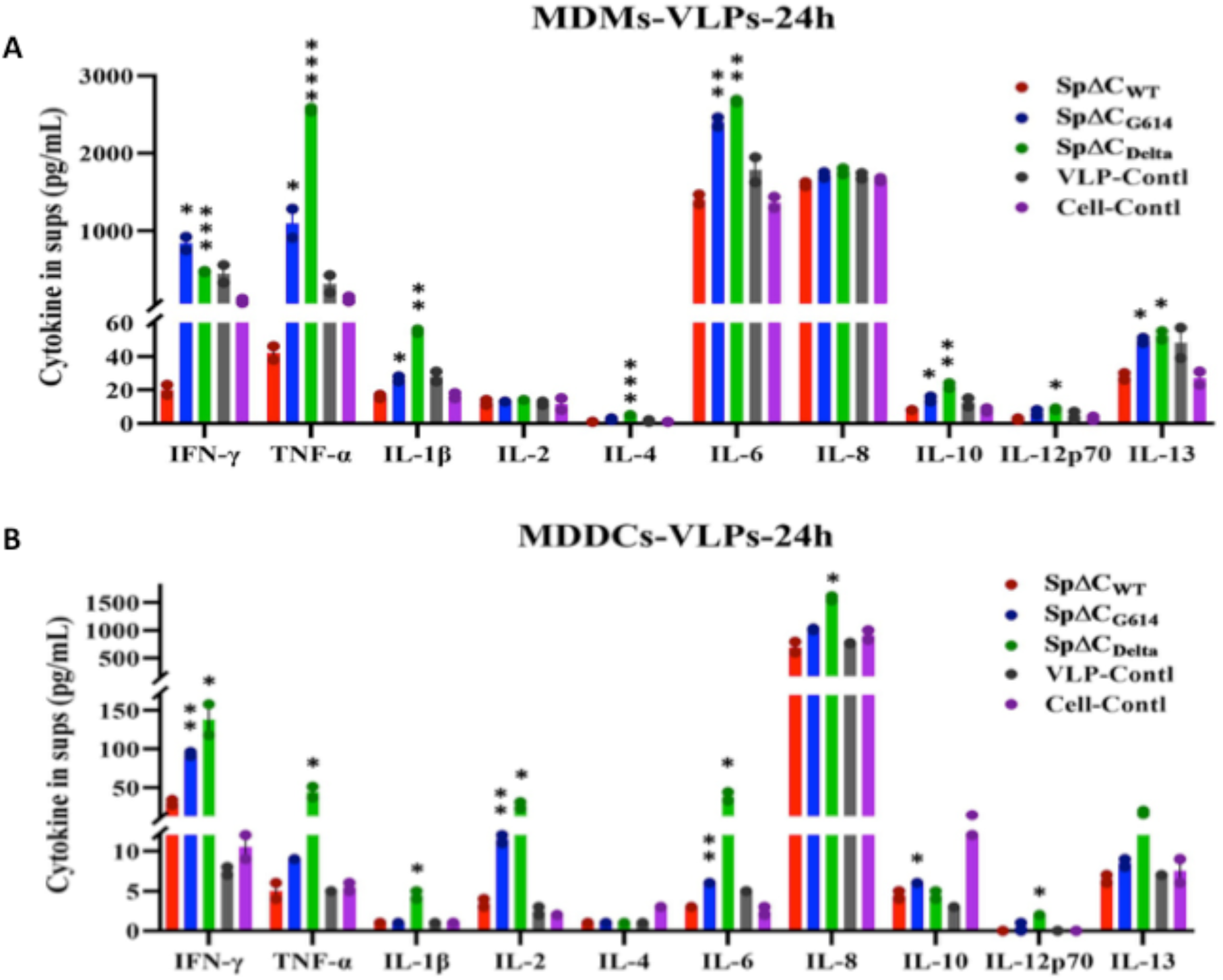
SpΔC_Delta_-PVLPs stimulated proinflammatory cytokines release in human MDM and MDDCs. Human MDMs (A) or MDDCs (B) were treated with equal amount of SPΔC_wt_, SPΔC_G614_ or SPΔC_Delta_ –PVLPs (adjusted by p24). Gag-VLPs treated and non-treated MDMs or MDDCs were used as negative controls. After 24 hrs, the cell culture supernatants were collected and different cytokines, including IFN-γ, IL-1β, IL-2, IL-4, IL-6, IL-8, IL-10, IL-12p70, IL-13, TNF-α were measured by MSD immunoassay (Meso Scale Discovery). The results are the mean values ± standard deviations (SD) of two biologic replicates. Statistical significance was determined using unpaired t test, and significant p values are represented with asterisks (*P ≤0.05; **P ≤ 0.01; ***P≤0.001; **** P≤0.0001). No significant (ns) was not shown.

Surprisingly, in MDDCs, SPΔC_Delta_-PVLP stimulation resulted in a significant increase in most pro-inflammation cytokines we tested, including IFN-γ, TNF-α, IL-1β, IL-2, IL-6, IL-8, and IL-12p70 (Fig. 5B). Among them, IL-6, TNF-α and IL-2 were the most increased cytokines (8∼13-fold), followed by IFN-γ and IL-1β (5∼6-fold). The levels of IFN-γ, IL-2, and IL-6 in the supernatans of SPΔC_G614_-PVLPs treated MDDCs were higher than that of SPΔC_WT_-PVLPs or control VLPs, also to a less extent. In contrast, IL-10 production appears to be negatively regulated by all SPΔC-PVLPs, including the control VLPs. Altogether, the above results suggested that SPΔC_Delta_ could induce remarkably higher levels of most proinflammatory cytokines tested and small differentce in some anti-inflammatory cytokines than VLP-SPΔC_G614_ or SPΔC_wt_ in human MDMs and MDDCs.

## DISCUSSION

The SARS-CoV-2 Delta variant has higher transmissibility and thus has become the predominant strain worldwide in 2021(Li et al., 2021). It is important to understand the mechanisms of the increased transmissibility and cytokine release triggered by this variant. In this study, we investigated the cleavage and maturation efficiency of the Delta variant Spike protein (SPΔC_Delta_) during pseudovirus assembly and its impact on cell-free pseudovirus infection and cell-cell fusion activities. The results demonstrated enhanced cleavage and maturation of SPΔC_Delta_ in the produced viral particles. Additionally, the studies clearly showed that SPΔC_Delta_ mediated more efficient pseudovirus infection and mediated a significantly enhanced cell-cell fusion process. Furthermore, our analyses revealed that SP_Delta_-PVLPs had stronger effects on stimulating NF-κB and AP-1 signaling in THP1 cells and elevated the production of several essential proinflammatory cytokines by human MDMs and MDDCs, compared with SP_WT_-PVLPs.

SARS-CoV-2 transmission and pathogenesis require the polybasic cleavage site between S1 and S2 in its SP (Johnson et al., 2021; Peacock et al., 2021a). This furin protease cleavable site is critical for the maturation of SARS-CoV-2 and its biological functions (Boson et al., 2021; Hoffmann et al., 2018; Hoffmann et al., 2020a). In an attempt to understand the mechanisms by which the Delta variant is more infectious than other variants, our study revealed that SPΔC_Delta_ was significantly more efficient in the processing of the precursor S into S1 and S2 compared with SPΔC_WT_ and SPΔC_G614_ in both the cells and PVPs. In the PVPs, the majority of SPΔC_Delta_ was presented as mature forms (S1 and S2), while some portions of the precursor S of SPΔC_WT_ and SPΔC_G614_ were still present in the PVPs (Fig. 1C, right panel). To further investigate the importance of the furin cleavage site for the enhanced cleavage of the Delta variant, we produced SPΔC_Delta_-PVPs packaged in the presence or absence of the furin protease inhibitor CMK. As expected, CMK significantly inhibited the cleavage of the spike protein of the Delta variant.

Among the mutations in the Delta variant spike protein, an amino acid proline (P681) at the N-terminus of the polybasic cleavage site (RRAR) was changed to arginine (R), known as the P681R mutation (Saito et al., 2021). The P681R mutation is of great importance because it is part of a proteolytic cleavage site for furin and furin-like proteases. The P618R mutation clearly plays a critical role in the SP of the Delta variant to abrogate host *O-*glycosylation (Zhang et al., 2021b). To further investigate whether this altered polybasic cleavage site (RRRAR) is required for effective furin cleavage of SARS-CoV-2 SP, we reverted the arginine (R) of 681 and asparagine (N) of 950 back to the original proline (P) and aspartic acid (D) on SP of the Delta variant (SPΔC_Delta-PD_). The WB results showed that the cleavage of the SPΔC_Delta-PD_-PVPs was comparable to that of SPΔC_WT_-, SPΔC_G614_- or SPΔC_Delta_-PVPs produced in the presence of CMK. Consistent with other reports (Peacock et al., 2021b; Saito et al., 2021), our results further demonstrated that the P618R mutation in SPΔC_Delta_ is essential for the enhanced furin cleavage of Delta variant SP. Meanwhile, we also observed that S1 of SPΔC_Delta_ appeared to migrate faster than S1 of SPΔC_WT_ and SPΔC_G614_. Although the mechanism is currently unknown, it could be related to the possible altered glycosylation content of SPΔC_Delta_ because a previous study revealed that P681R could circumvent host *O*-glycosylation (Zhang et al., 2021b). In addition, multiple amino acid mutations and deletions present in S1 of Delta SP may also partially contribute to this alteration. More detailed studies are required to analyze the possible underlying mechanism(s).

In this study, we also observed that SPΔC_Delta_ enhanced cell-free pseudovirus infection in A549_ACE2+_ and Calu-3 cell lines and macrophages, indicating that SPΔC_Delta_ is an important factor contributing to the increased infectiousness of the Delta variant. Additionally, it was observed that the infection mediated by SPΔC_Delta_-pseudovirus was only slightly higher than that mediated by SPΔC_G614_-pseudovirus, suggesting that the G614 mutation present in SP_Delta_ may be one of the main driving forces for the increased infectivity of the Delta virus (Daniloski et al., 2020; Zhang et al., 2020). This finding is in agreement with previous studies showing that the D614G mutation in SARS-CoV-2 SP contributes immensely to virus infectivity and replication (Daniloski et al., 2020; Zhang et al., 2020). It should be noted that our results were based on single-cycle SPΔC-pseudovirus replication; thus, the infection advantage of the native SARS-CoV-2 Delta variant needs further investigation.

It is well known that spike protein expressed at the surface of infected cells is sufficient to generate fusion with neighboring cells. Here, we further showed that significantly enhanced syncytia formation was observed when SPΔC_Delta_-expressing 293T or A549 cells were cocultured with A549_ACE2_ cells. This observation raised potential importance in terms of the SARS-CoV-2 Delta variant’s virulence since cell-to-cell fusion may provide another efficient method of viral dispersal in the host, thus indicating its stronger transmission among the population, as described previously (Michael Rajah et al., 2021). This finding is in agreement with recent reports that B.1.617.2 SP mediates highly efficient syncytia formation compared with wild-type SP (Michael Rajah et al., 2021; Mlcochova et al., 2021; Planas et al., 2021; Zhang et al., 2021a). Moreover, this efficient cell-to-cell transmission ability of SPΔC_Delta_ may enhance its resistance to host immune responses, such as antibody-mediated neutralization (Mlcochova et al., 2021; Planas et al., 2021). It is worth noting that we did not investigate the impact of TMPRSS2 on the cell-cell fusion process. For SARS-CoV-2, cleavage of S by furin at the S1/S2 site is required for subsequent cleavage by TMPRSS2 at the S2’ site. Previous studies have demonstrated that TMPRSS2 could enhance the infectivity and fusogenic activity of different coronaviruses, including SARS-CoV-2 (Buchrieser et al., 2020; Glowacka et al., 2011; Kleine-Weber et al., 2018; Matsuyama et al., 2010). Future investigations into the role of TMPRSS2 in SP Delta-induced syncytia formation and infection will provide a better understanding of the persistence, dissemination, and immune or inflammatory responses of Delta variants.

The severity of COVID-19 is highly correlated with dysregulated and excessive release of proinflammatory cytokines (Huang et al., 2020a). Hence, we also tested whether macrophages or dendritic cells act as major modulators of the immune response by producing a large amount of cytokines and chemokines to recruit immune cells and presenting antigens to them. The engagement of the spike protein of SARS-CoV-2 with the receptor ACE2 on THP1-derived macrophages is reported to initiate signaling pathways and activate the production of proinflammatory cytokines, including IL-6, TNF-α, and MIP1a (Pantazi et al., 2021). Here, we showed that NFκB and AP1 signaling pathway activities were also enhanced by SPΔC_Delta_ compared with SPΔC_WT_ in THP1 cells and THP1-derived macrophages, suggesting that SPΔC_Delta_ might promote the inflammatory status of these cells. Similarly, the SP of SARS-CoV-1 has been discovered to activate NF-κB and stimulate the release of IL-6 and TNF-α (Wang et al., 2007).

A previous study reported that high plasma levels of TNFα, IL-1, IL-6, IL-8 and other inflammatory mediators were found in severe COVID-19 patients, and the serum IL-6, IL-8 and TNF-α levels were strong and independent predictors of disease progression, severity and death. (Del Valle et al., 2020; Huang et al., 2020a; Santa Cruz et al., 2021). Interestingly, we found that SPΔC_Delta_ significantly enhanced the expression of several proinflammatory cytokines (TNF-α, IL-1β, and IL-6) in both MDMs and MDDCs (Fig. 5). Especially for MDDCs, increased levels of other proinflammatory cytokines, including IFN-γ, IL-2, IL-8, and IL-12p70 were also detected (Fig. 5). However, SPΔC_wt_ only exhibited induction of IFN-γ in MDDCs, but did not show any effect on other cytokine production of MDMs or MDDCs. This is agree with previous study that revealed, upon SARS-CoV-2 infection, neither macrophage, nor dendritic cells produce the pro-inflammatory cytokines (Niles et al., 2021). Mutations in Delta SP seem to be the key points that cause the differential expression of cytokines. Given the fact that IFN-γ, TNF-α, IL-1β and IL-12 are T-helper-1 (Th1) cytokines, it also suggests that the Th1/Th2 balance has further shifted to Th1 dominance after stimulation with SPΔC_Delta_-PVLP. Along with proinflammatory cytokines, three anti-inflammatory cytokines (IL-4, IL-10 and IL-13) were also increased in SPΔC_Delta_-PVLP treated macrophages compared with SPΔC_WT_-PVLP_-_treated macrophages. Consistently, higher secretion of T-helper-2 (Th2) cytokines such as IL-4 and IL-10 has been reported in ICU patients than in non-ICU patients (Huang et al., 2020a). Their functions are to suppress both inflammation and the TH1 cellular response, indicating that the balances between pro- and anti-inflammation, as well as the balances between TH1 and TH2 cellular responses existing in patients, are important for the clinical outcomes of COVID-19 therapy. However, the lower IL-10 level in all PVLPs treated MDDCs is surprising and the reason of this is unclear. In conclusion, SPΔC_Delta_-treated macrophages and DCs are in a higher inflammatory state and in a Th1-dominant Th1/Th2 balance.

Overall, we demonstrated that the SARS-CoV-2 Delta variant spike protein exhibited enhanced cleavage and maturation, which may play an important role in viral infection and cell-cell transmission. Furthermore, we revealed that SP_Delta_ had stronger effects on stimulating NF-κB and AP-1 signaling in monocytes and the release of proinflammatory cytokines from human macrophages and dendritic cells. All of these studies provide strong evidence to support the important role of Delta SP during virus infection, transmission and pathogenesis.

## MATERIALS AND METHODS

### Plasmid constructs

The SARS-CoV-2 SP protein-expressing plasmids (pCAGGS-nCoVSPΔC and pCAGGS-nCoVSPΔC_G614_) were described previously (Ao et al., 2021a). The gene encoding SPΔC_Delta_ or SPΔC_Delta-PD_ was synthesized (Genescript) and cloned into the pCAGGS plasmid, and each mutation was confirmed by sequencing. pEF1-SPΔCwt, pEF1-SPΔC_G614_ or pEF1-SPΔC_Delta_ was constructed by inserting the cDNA encoding SPΔCwt, SPΔC_G614_ or SPΔC_Delta_ through the *BamHI* and *NheI* sites into the pEF1-pcs-puro vector (Ao et al., 2008). The HIV RT/IN/Env trideleted proviral plasmid containing the Gaussia luciferase gene (ΔRI/E/Gluc) and the helper packaging plasmid pCMVΔ8.2 encoding the HIV Gag-Pol plasmids have been described previously (Ao et al., 2016; Zhang et al., 2016).

### Cell culture, antibodies and chemicals

Human embryonic kidney cells (HEK293T), human lung (carcinoma) cells (A549), A549_ACE2_, Calu-3 cells and THP1-sensor cells were cultured in Dulbecco’s modified Eagle’s medium or RPMI 1640 medium supplemented with 10% fetal bovine serum (F.B.S.) and 1% penicillin/streptomycin. To obtain human MDMs or MDDCs, human peripheral blood mononuclear cells (hPBMCs) from healthy donors were collected by sedimentation on a Ficoll (Lymphoprep; Axis-Shield) gradient, adherent to 24-well plates for 2 hrs, and then treated with macrophage colony stimulator (M-CSF) or granulocyte-macrophage-stimulating factor (GM-CSF) and IL-4 (R&D system) for 7 days.

The THP1-NF-κB-Luc and THP1-AP-1-Luc sensor cell lines were described previously (Ao et al., 2021b). To obtain THP1-derived macrophages, THP-1-NF-κB-Luc and THP1-AP-1-Luc sensor cell lines were treated with phorbol 12-myristate 13-acetate (PMA) (200 ng/mL) for 3 days followed by 2 days of rest, as previously described (Starr et al., 2018). A549-expressing human ACE2 (A549_ACE2_) cells were generated by transducing A549 cells with the ACE2-expressing lentiviral vector (pLenti-C-mGFP-ACE2) (Origene, Cat# PS100093) and then selected with puromycin according to the manufacturer’s procedure.

The rabbit polyclonal antibody against SARS-CoV-2 SP/RBD (Cat# 40592-T62) or human SARS-CoV-2 S-NTD antibody (E-AB-V1030) was obtained from Sino Biological or Elabscience. Mouse monoclonal antibody (1A9) against SARS-CoV-2 SP-S2 (Cat# ab273433) was obtained from Abcam. Anti-HIVp24 monoclonal antibody was described previously (Ao et al., 2007; Qiu et al., 2011). Anti-human ACE2 antibody (sc-390851) was obtained from Santa Cruz Biotechnology Inc. Furin inhibitor I, a peptidyl chloromethylketone (CMK) (Cat# 344930), was obtained from Millipore Sigma.

### Virus production and infection experiments

SARS-CoV-2 SPΔC pseudotyped viruses (CoV-2-SPΔC-PVs, CoV-2-SPΔC_G614_-PVs and CoV-2-SPΔC_Delta_-PVs) or pseudotyped virus-like particles (VLPs) were produced by transfecting HEK293T cells with pCAGGS-SPΔC_WT_, pCAGGS-SPΔC_G614_ or pCAGGS-SPΔC_Delta_ and pCMVΔ8.2 with or without a Gluc-expressing HIV vector ΔRI/E/Gluc (Ao et al., 2021a). After 48 hrs of transfection, cell culture supernatants were collected, and VPs or VLPs were purified from the supernatant by ultracentrifugation (32,000 rpm) for 2 hrs. The pelleted VPs or VLPs were resuspended in RPMI medium, and virus titers were quantified by HIV-1 p24 amounts using an HIV-1 p24 ELISA.

To measure the infection ability of SARS-CoV-2 SPΔC pseudotyped VPs, equal amounts of each SPΔC-PVs stock (as adjusted by p24 levels) were used to infect A549_ACE2_, Calu-3 cells, human MDMs or MDDCs. After different time intervals (24, 48 and 72 hrs), the supernatants were collected, and the viral infection levels were monitored by measuring Gaussia luciferase (Gluc) activity. Briefly, 50 µl of coelenterazine substrate (Nanolight Technology) was added to 10 µl of supernatant, mixed well and read in a luminometer (Promega, U.S.A.).

To evaluate the effects of various SCoV-2 SPΔC-VLPs on the NF-κB and AP-1 signaling pathways, the same amount of each SPΔC-pseudotyped VLP stock (10 ng, as adjusted by the p24 levels) was directly added to THP1-NF-κB-Luc or THP1-AP1-Luc sensor cells. After 6 hrs, the cells were collected and subjected to luciferase assay as described previously (Ao et al., 2021b). To test the effect of different SPΔC-VPs on cytokine production in MDMs and MDDCs, the same amount of each SPΔC-VP stock (20 ng, as adjusted by the p24 levels) was added to MDMs and MDDCs, and the supernatants were collected after 24 hrs. The cytokine (IFN-γ, IL-1β, IL-2, IL-4, IL-6, IL-8, IL-10, IL-12p70, IL-13, TNF-α) levels in the supernatants were measured using the MSD V-PLEX proinflammatory Panel 1 (human) Kit (Mesoscale Discovery, USA, Cat# K15049D-1) following the manufacturer’s procedure.

### Generation of different SPΔC-expressing A549 stable cell lines

Production of lentiviral vectors expressing different SPΔC: 293T cells were cotransfected with pEF1-SPΔC_wt_, pEF1-SPΔC_G614_ or pEF1-SPΔC_Delta_ with packaging plasmid Δ8.2 and VSV-G expressing plasmid. Forty-eight hours posttransfection, each lentiviral vector particle in the supernatant was collected. Then, each produced lentiviral particle was used to transduce A549 cells, and the transduced cells were selected with puromycin for one week. SPΔCwt/mutant expression in the different transduced A549 cells was evaluated by WB using an anti-RBD antibody.

### Immunofluorescence assay

293T cells transfected with various SARS-CoV-2 SPΔC-expressing plasmids were grown on glass coverslips (12 mm^2^) in a 24-well plate. After 48 hrs, cells on the coverslip were fixed in 4% paraformaldehyde for 5 minutes and permeabilized with 0.2% Triton X-100 in PBS. The cells were then incubated with primary antibodies against the N-terminal domain of SARS-CoV-2 SP followed by the corresponding FITC-conjugated secondary antibodies. The cells were viewed under a computerized Axiovert 200 fluorescence microscope (ZEISS).).

### Syncytium formation assay

293T cells were transfected with pCAGGS-SPΔC_WT_, SPΔC_G614_, SPΔC_Delta_ or SPΔC_DeltaPD_ plasmids using Lipofectamine 2000. After 24 hrs, the cells were washed, resuspended and mixed with A549_ACE2_ cells at a 1:3 ratio and plated into 48-well plates. For syncytium formation of the stable cell line, A549-SPΔC_WT_ or A549-SPΔC_Delta_ cells were detached with 0.05% trypsin and mixed with A549 or A549_ACE2_ cells. At different time points, syncytium formation was observed, counted and imaged by bright-field microscopy (Axiovert 200, ZEISS).

### Statistics

Statistical analysis of cytokine levels, including the results of GLuc assay, Luciferase assay, and various cytokine/chemokines assay, were performed using the unpaired t-test (considered significant at P≥0.05) by GraphPad Prism 9 software.

## ACKNOWLEDGEMENTS

We thank Dr. Darwyn Kobasa for providing the Calu-3 cell lines and technique supports. Titus Olukitibi is a recipient of the University Manitoba Graduate scholarship. This work was supported by Canadian 2019 Novel Coronavirus (COVID-19) Rapid Research Funding (OV5-170710) by Canadian Institute of Health Research (CIHR) and Research Manitoba, and CIHR COVID-19 Variant Supplement grant (VS1-175520) to X-J.Y. This work was also supported by the Manitoba Research Chair Award from the Research Manitoba (RM) to to X-J.Y.

## AUTHOR CONTRIBUTIONS

Experimental design, X. Y, Z.A. and M.J.O; Investigation, Z.A., M.J.O., and O.T.A; Writing-Original Draft Preparation, Z.A. and M. J. O. and O.T.A; Review, Z.A. M. J. O. and X.Y. Supervision, X.Y.

